# Modulating neutrophils leads to changes in CNS output

**DOI:** 10.1101/2022.02.10.479966

**Authors:** A.P. Coulibaly, I. Salken, J.J. Provencio

## Abstract

It has become increasingly evident that immune cells play a role in modulating brain function. Neutrophils are the most abundant innate immune cell in mammals. Until recently, our understanding of the physiological role of these cells was limited to infection and cancer. Recently, it has emerged that neutrophils play critical roles in modulating tissue health and physiology. There has been evidence that neutrophils populate the rodent meninges. As such, we hypothesized that like other meningeal immune cells, neutrophils play a neuromodulatory role. Using a depletion paradigm, anti-Ly6G antibody injection, we demonstrate that the loss of neutrophils leads to changes in cognitive flexibility and social interactions in mice. Furthermore, the depletion paradigm led to a 50% decrease in the number of neutrophils in the meninges, which in turn led to the proliferation of resident immature neutrophils within the meninges. Finally, our data show that this immature population reenters the cell cycle only in response to specific physiological stimuli (neutrophil depletion, and stress). In conclusion, this study demonstrates that there are resident neutrophils within the meninges and that elucidating their function in the nervous system will deepen our understanding of immune regulation of brain function.

## Introduction

Recent evidence demonstrates that the meninges act as a source of immune cells for the nervous system ^1^. Indeed, the meninges have been shown to be a reservoir of immune cells, including myeloid cells ^2–5^ and lymphocytes ^6,7^. There is mounting evidence for the role of meningeal-based immune cells in the regulation of the behavior ^8^. Furthermore, the spinal meninges also contain immune cells that may play a role in the spinal response to injury and disease ^9^. We believe that understanding the role of meningeal immune cells under normal condition is critical to better understanding of non-neuronal influences on behavior.

Myeloid cells are a large subset of the innate immune system. In the periphery, they make up the early response system to pathogenic challenges. Some are critical to the removal of infectious agents^10^. Neutrophils are the most abundant myeloid cell and their activation has been linked to immune modulation, tissue damage and healing after injury ^11–13^. Our laboratory has shown that neutrophil function in the meninges is critical for the development of cognitive deficits after subarachnoid hemorrhage (SAH)^14^. As expected, most neutrophils infiltrating the meninges after SAH are blood derived, although we found the presence of neutrophils in the meninges prior to the injury suggesting a meningeal neutrophil population. This is supported by characterization of meninges in normal mice where a large proportion of meningeal immune cells are neutrophils ^15,16^. Because neutrophils are typically considered short-lived non-tissue resident cells, their presence in the normal meninges is unexpected and their role is not understood.

In the present study, we demonstrate that manipulating neutrophil levels within mice can lead to changes in behavioral output. Depletion of neutrophils, systemically, significantly impacts neutrophil dynamics within the meninges. We further demonstrate that the neutrophil population in the meninges is heterogenous including mature, activated, infiltrating, senescent, and immature neutrophils. This immature population appears to be a progenitor subset that can divide and partially populate the meninges upon stimulation. Furthermore, we show that mature neutrophils, derived from progenitors in the meninges, enter the skull marrow space, possibly as a site of degradation. The result of this work provides evidence of the meninges as a local source of neutrophils, both marginated and new cells, for the central nervous system.

## Results

### Neutrophil depletion alters cognitive flexibility and social interactions in mice

Our results demonstrate that sustained neutrophil depletion, using serial injections of anti-Ly6G (every 2 days), leads to abnormal mouse behavior. Specifically, changes were observed in the Barnes maze (Figure 1A), novel object recognition (Figure 1B), and social interaction (Figure 1C) which will be outlined below.

**Figure 1:**
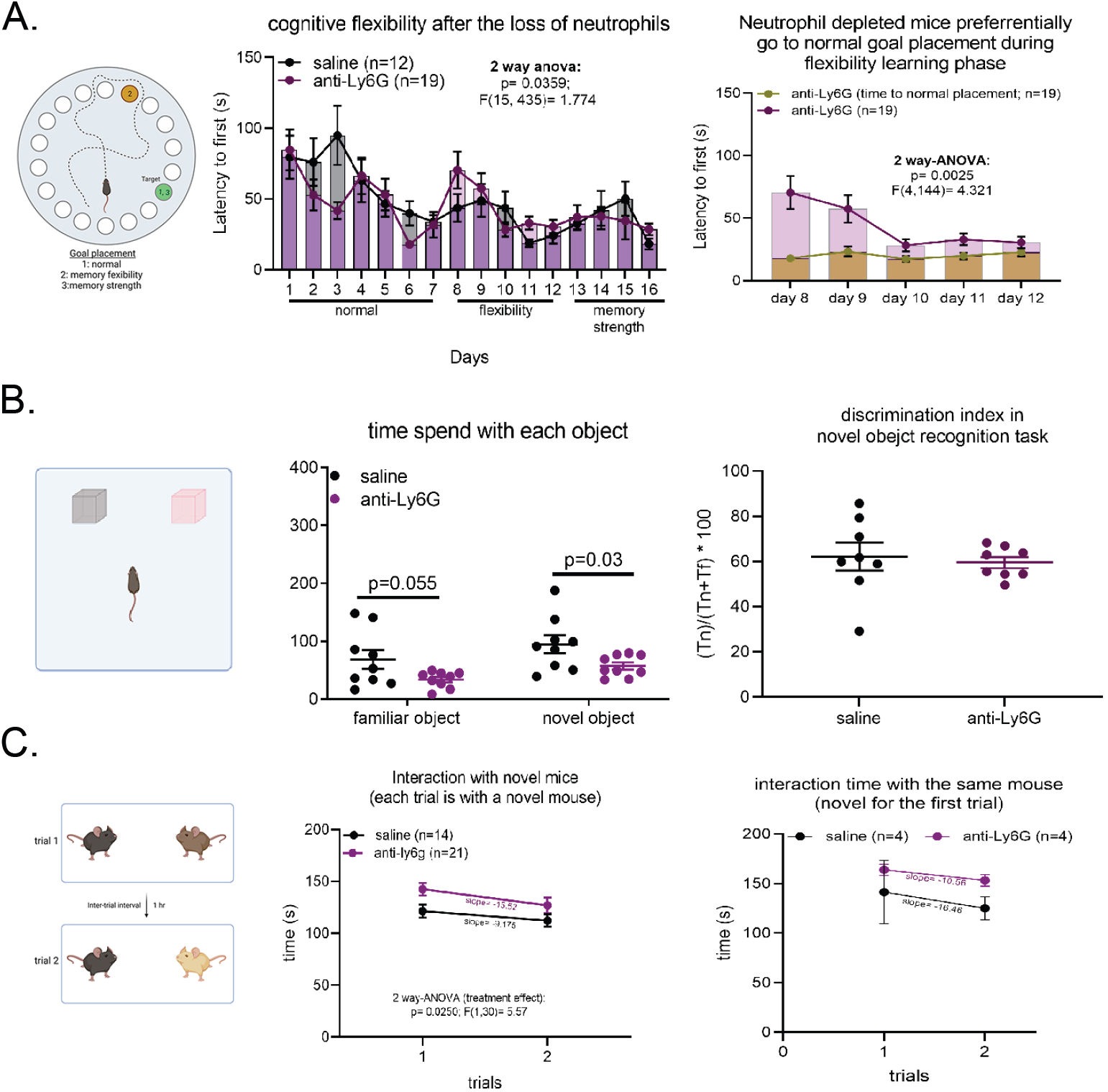
Neutrophil depletion leads to behavior changes in mice. A) Mice with chronic neutrophil depletion show a deficit in reversal learning (cognitive flexibility) in the Barnes maze task, with depleted mice performing poorly when the goal box is moved. Post-hoc analysis shows a preference to the original placement of the goal box during the flexibility portion of the experiment. B) In a novel object recognition task, mice without neutrophils spend significantly less time with both the familiar and novel object during exploration. However, no difference is observed between in the discrimination index of both groups. C) In a social interaction task, mice lacking neutrophils spend significantly more time interacting with the first novel mouse they encounter. However, this interaction decreases when exposed to second unfamiliar mouse in similar fashion to that of the control. When exposed to the same mouse, the neutrophil depleted mouse spend more time with the now “familiar” mouse than control. Statistical significance, p>0.05; 2 way-ANOVA used in A) and C), and student’s T-test used for both B).

Using a modification of the Barnes maze as a measure of cognitive flexibility, mice without neutrophils perform statistically better than saline injected mice during the normal learning phase of the task (Figure 1A). However, when the goal box is moved (to test reversal learning or cognitive flexibility), these mice are less able to adapt to the new goal box placement. Indeed, neutrophil-depleted mice preferentially visit the site of the original goal box before going to the new location. After the reversal learning phase, when the goal box is placed in its original location, neutrophil-depleted mice do as well than control mice (similar to their performance during the normal learning phase of the task). These data suggest that the loss of neutrophils is affecting spatial memory and learning which make up the ability to be cognitively flexible.

The finding of decreased cognitive flexibility were replicated using a puzzle box task (Supplemental figure 1). This test shows that changing the obstacle at the entrance of the safety box significantly affects neutrophil-depleted mice’s performance in the task.

Informed by the changes in cognitive flexibility, we tested whether hippocampal mediated memory acquisition and retention are affected by performing a novel object recognition (NOR) task. Neutrophil-depleted mice spend less total time with both objects when compared with saline injected mice, although the percentage of time they spend with each object (the discrimination index) is the same (Figure 1B). These data suggest that although exploration times differ, neutrophil-depleted mice are able to accurately differentiate between a novel and familiar object.

To determine whether the lower interaction is specific to inanimate objects, a social interaction experiment was performed, where mice were introduced in consecutive trials to either two different (novel) mice or the same mouse twice. Unlike in the novel object recognition, neutrophil-depleted mice spend more time with the non-familiar mouse than control (Figure 1C). Indeed, when exposed to two different novel mice, the neutrophil-depleted and control mice interact equally with each mouse (Supplemental figure 2). On the other hand, when exposed to the same mouse twice, the control mouse spends less time with the non-novel mouse in the second trial but the neutrophil-depleted mouse treats the test mouse as if it were a novel mouse (spending the same amount of time with the mouse as it did when the test mouse was novel) (supplemental figure 2).

### Neutrophil population in the meninges is heterogenous under normal conditions

Recent studies have demonstrated that immune cells in the meninges influence behavioral outcome ^17,18^. Because neutrophils populate the meninges, it is important to determine if that population is affected in our depletion paradigm. As such, we first characterized whether intraperitoneal administration of anti-Ly6G leads to changes in the neutrophil composition of the meninges. Indeed, this paradigm leads to a 50% decrease in the neutrophil in the meninges (Supplemental figure 3). Therefore, to truly understand the function of these neutrophils in the meninges, we took a deeper look at the neutrophil population within the meninges.

Our analysis demonstrates that neutrophils populate the meninges under healthy conditions (Figure 2A). They are found in both the meningeal capillary beds and the fibrous parenchyma in all three meningeal layers (Figure 2A). A proportion of neutrophils in the mouse meninges exhibit circular nuclei, suggesting an immature profile. Using an imaging flow cytometry method, of dissociated meninges, we identified cells with mature, hyper-segmented nuclei, and immature, banded or circular nuclei (Figure 2B). Using spectral flow cytometry, we identified both (Figure 2C), mature (Ly6G^hi^) and immature (Ly6G^int^, SCA1, CD44, CD34) neutrophils. Among the mature neutrophils, there are activated (Ly6G^hi^, CD83), infiltrating (Ly6G^hi^, CD62L), and senescent (CXCR4, CD62L, Ly6B) cells.

**Figure 2:**
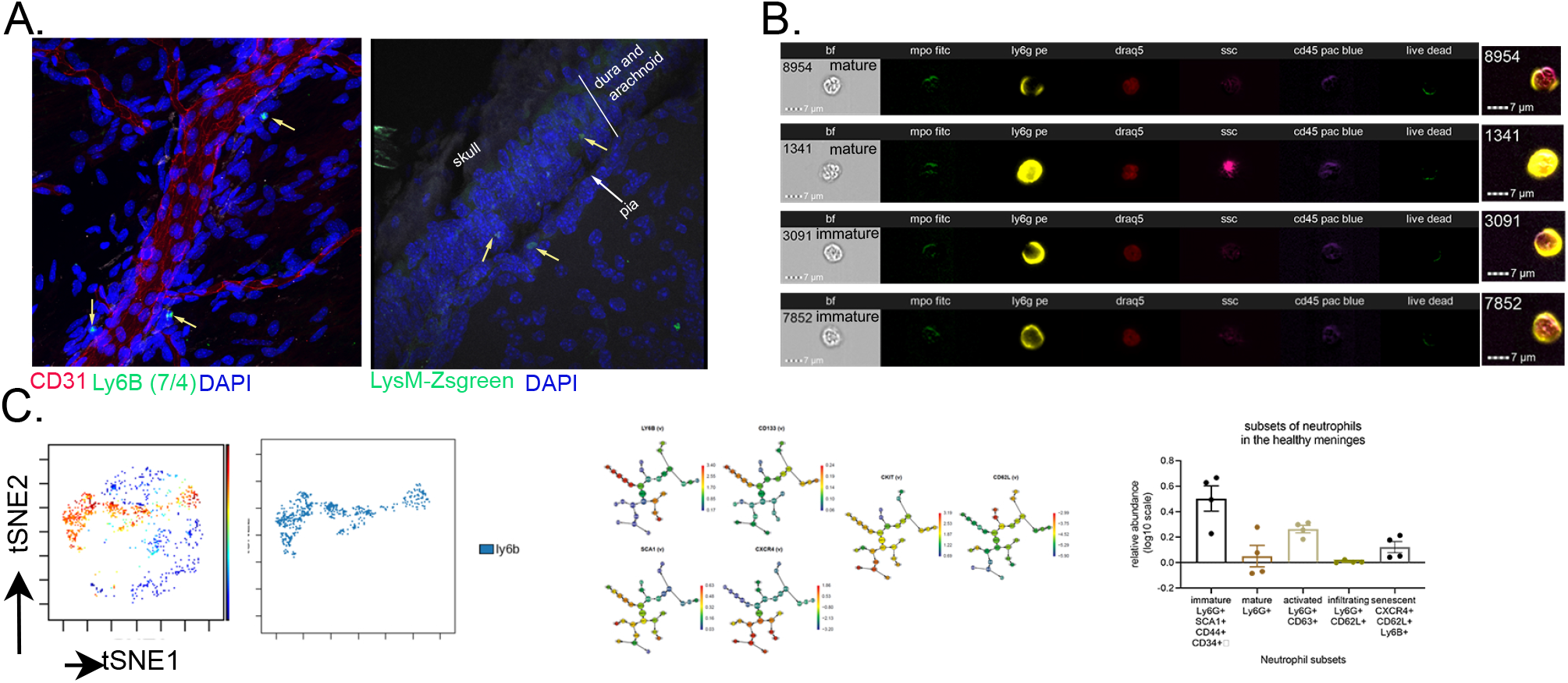
A heterogenous neutrophil population is found in the meninges of the naïve mouse. A) Confocal micrograph of the naïve mouse meninges shows the presence of neutrophils in the meninges and adjacent blood vessels. Closer characterization shows neutrophils located within all meningeal layers, dura, arachnoid and pia. B) Imaging flow cytometry (ImageStream) shows the presence of immature and mature neutrophils within the meninges of the mouse. C) Spectral flow analysis combined with Cytobank demonstrate the presence of immature, mature, activated, infiltrating, and senescent neutrophils within the naïve mouse meninges.

To understand the dynamics of these cells in the meninges, we investigated whether the immature population is resident to the meninges and, if so, can be induced to proliferate in situ. In the peripheral environment, the release of immature neutrophils from the bone marrow is associated with infection^19^. Since our mice were healthy, we presumed that the immature cells observed were not due to the release from the bone marrow. Recently, multiple sources have demonstrated that the spleen, unlike other organ, has its own reservoir of neutrophils, including an immature/progenitor population ^20,21^. This population is critical to splenic response to fungal infection and the maintenance of cells in the spleen marginal zone^11,20^. To determine whether the meninges had its own reservoir (i.e. resident) of neutrophils, we used a radiation-induced chimeric model in the LysM-tdTomato mouse, in which all myeloid cells including neutrophils are tdTomato^+^. These mice are exposed to two doses of gamma irradiation (350 rad for the first dose, and 950 rad for the second) without head shielding to deplete neutrophils in the head including the skull bone marrow. The doses are separated by 48 hours, then the recipient mice are transplanted with bone marrow from a UBC-GFP donor mouse, in which all cells are GFP^+^. As such, the resulting chimeric mice, with full bone marrow reconstitution exhibit GFP^+^ bone marrow-derived immune cells. We then characterized, in the irradiated mouse, the percentage of neutrophils in the bone marrow, blood and meninges from the donor (GFP^+^) or recipient mouse (survivor neutrophils that remain tdTomato^+^) using flow cytometry 6 weeks after the second dose of radiation.

In all tissue types analyzed, most neutrophils were GFP^+^, suggesting that they originate from the transplanted bone marrow. Specifically, 97% of all bone marrow cells are GFP^+^ (3 out of the 4 mice showed no tdTomato^+^ neutrophils), 100% of the circulating neutrophils in the blood are GFP^+^, but only 87% of the neutrophils in the meninges are GFP^+^ (Figure 3A). 13% of the neutrophils in the meninges are either from the immature neutrophil subset who survived irradiation or are long-lived (6 weeks) mature neutrophils from before irradiation. It is well established that neutrophils are short-lived ^22^. So, we explore the possibility of a slow dividing, radioresistant, immature neutrophil population that will give rise to the tdTomato^+^ neutrophils present in the irradiated mouse meninges. Unlike the blood and bone marrow, 1.5% of the tdTomato^+^ neutrophils in the meninges express the stemness factor Ckit, suggesting a proliferative potential (Figure 3B). This does not appear to be a reaction to the chimerism since under healthy conditions, 0.3% of meningeal neutrophils are Ckit^+^ (Figure 3B). The higher percentage in the chimeric mouse may be a result of fewer tdTomato neutrophils in the meninges. These data suggest that the meninges contain a neutrophil progenitor population that contributes to the neutrophil number within the meninges.

**Figure 3:**
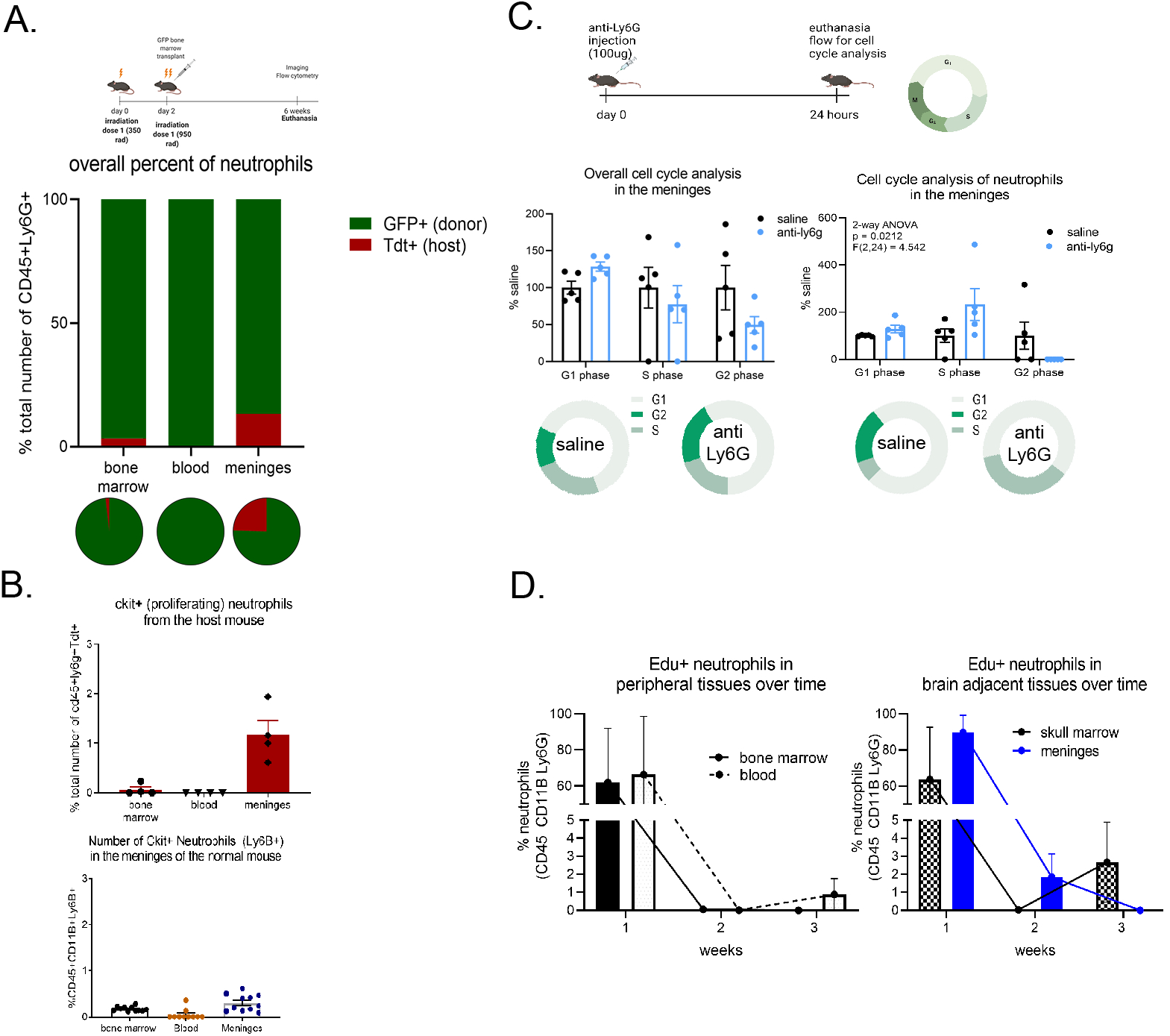
Resident immature neutrophils contribute to the neutrophil population within the naïve meninges. A) Using a full body irradiated bone marrow chimera model, with donor marrow (GFP) and recipient mouse (Tdtomato), we see the presence of long-lived (6 weeks) Tdtomato^+^ neutrophils (red) in the meninges of the mouse. B) Determination of stemness, using Ckit expression, showed that a portion of the Tdtomato neutrophils were Ckit^+^ after irradiation, demonstrating a proliferative potential. In addition, characterization of non-irradiated mice showed the presence of Ckit^+^ neutrophils in the mouse meninges. C) Immature neutrophils in the meninges re-enter the cell cycle after anti-Ly6G antibody neutrophil depletion. D) A single pulse EDU experiment shows the immature neutrophils in the meninges to be slow dividing.1 week after injection, EdU^+^ neutrophils were found in the bone marrow of long bones, blood, meninges, and skull marrow. 2 weeks after the injection, EdU^+^ were only present in the meninges. Finally, 3 weeks after the pulse injection of EDU, EDU^+^ neutrophils are only located in the skull marrow and the blood space.

To explore whether the progenitor neutrophil population in the meninges can be induced to proliferate under normal conditions, we employed the neutrophil depletion paradigm used in the behavioral experiments. Intraperitoneal injection of anti-Ly6G antibodies deplete neutrophils in the meninges (supplemental figure 3). Previous studies have demonstrated that anti-Ly6G depletion of neutrophils leads to the depletion of mature, fully-differentiated, neutrophils ^23^. Using flow cytometry staining for cell cycle analysis, we found that 24 hours after depletion there is a complete loss of neutrophils in the G2 phase, suggesting that these cells underwent mitosis to replace the mature neutrophils that were depleted (G1 phase). There is also an increase in the number of cells replicating their DNA, which is further evidence of a switch to proliferation in the immature neutrophil population. These results provide evidence that the immature neutrophils within the meninges can be stimulated to proliferate.

We next investigated whether immature neutrophils remain in the meninges for their entire life cycle. Under normal conditions, neutrophils are produced in the bone marrow and degraded in the bone marrow^24,25^, liver^26,27^, and lungs^28,29^. To investigate whether neutrophils leave the meninges at the end of their journey, we used a single EdU injection to track the location of newly made neutrophils in the blood, long bone marrow, skull marrow, and meninges (Figure 2D). One week after injection, most neutrophils at all these locations (on average 60-65% in the blood, skull and bone marrow; 90% in the meninges) were EdU^+^ suggesting that proliferating cells taking up EdU had divided to renew neutrophil populations. Two weeks after the injection, only the meninges contained EdU^+^ neutrophils (2%) suggesting that these neutrophils have remained in the meninges. At three weeks, no EdU^+^ neutrophils are found in the meninges, suggesting that the immature cells that had first incorporated EdU had undergone enough mitotic events to dilute the EdU concentration within the nucleus. Also, in the skull marrow, EdU^+^ cells were found at one week, were not found at 2 weeks, but recurred in the marrow at three weeks (3% of total neutrophils) suggesting a migration of EdU^+^ neutrophils from the meninges to the skull marrow space.

### Meningeal neutrophil numbers are tightly regulated

As demonstrated above, the deletion of neutrophils leads to behavioral changes in the mouse. Therefore, to determine whether specific CNS conditions can affect the immature neutrophil population in the meninges, we characterized the effect of enriched environment, acute stress, or injury on the number of immature neutrophils within the meninges. We identified these cells through their expression level of Ly6G. It is established that immature/progenitors of neutrophils have an intermediate expression of Ly6G while mature cells have high expression^30,31^. Our data show that environmental enrichment alone has little effect on the level of immature neutrophils (Ly6G^int^) in the meninges (Figure 4A). As expected, in both naïve and enriched environment, the depletion of neutrophils using anti-Ly6G antibodies leads to an increased number of immature neutrophils (no effect from the enrichment on this population; Figure 4A). To determine the effect of injury on the immature population, we looked at the levels of immature neutrophils in the meninges at 1 and 3 days after subarachnoid hemorrhage. Our laboratory has shown that subarachnoid hemorrhage leads to a significant increase of mature neutrophils in the meninges^14^. Surprisingly, our data show that the injury has no effect on the number of immature neutrophils in the meninges (Figure 4B). Interestingly, exposure to acute stress (restrain) leads to a marked increase in the number of immature neutrophils in the meninges (Figure 4C). These data suggest that the response of immature neutrophils in the meninges is context dependent.

**Figure 4:**
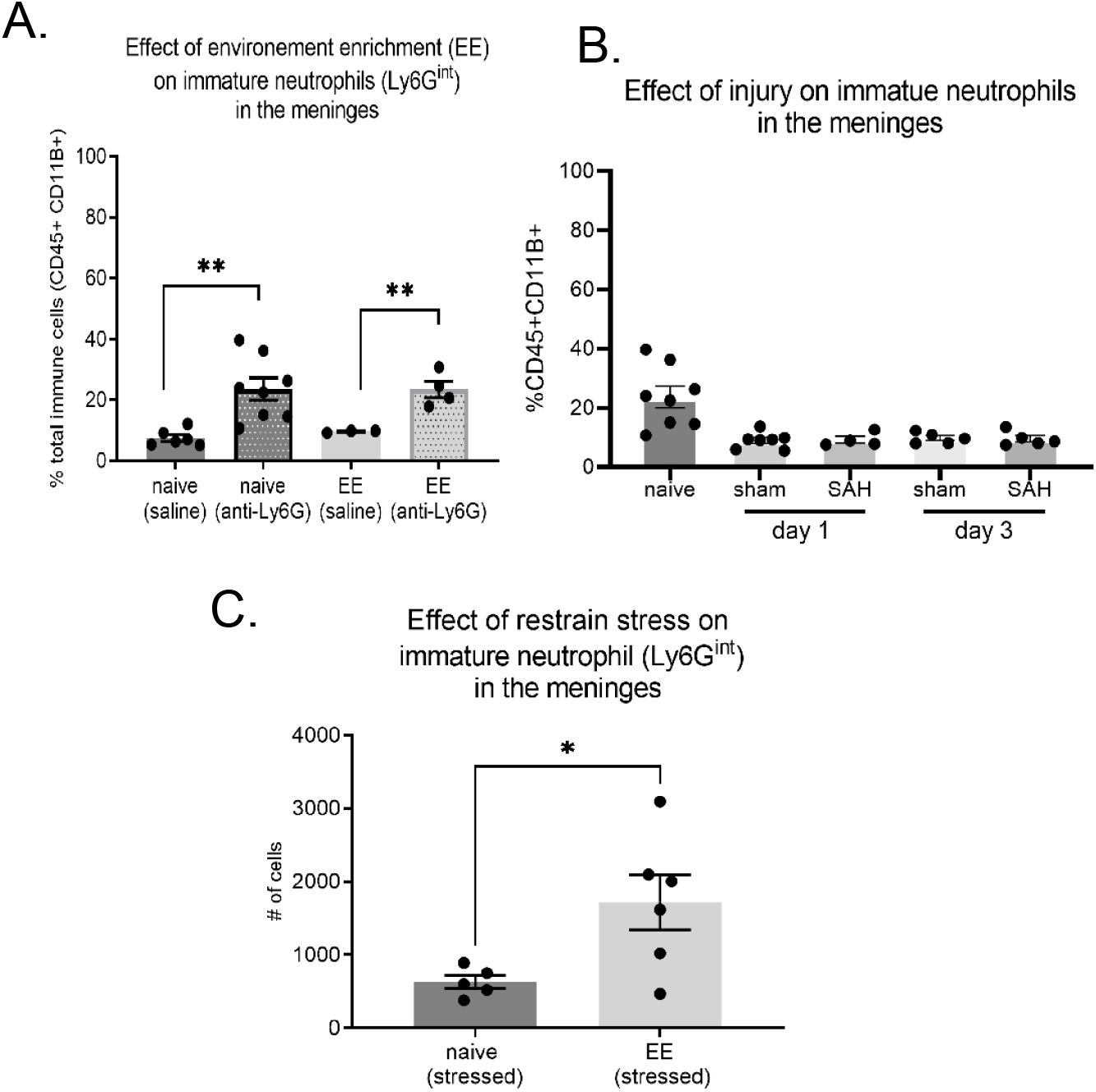
Immature neutrophils only divide under specific physiological conditions. A) Enriched environment does not further increase the rate of proliferation of immature neutrophils after anti-Ly6G depletion. Injury by subarachnoid hemorrhage does not affect the number of immature neutrophils in the meninges. B) Conversely, a continuous restraint stress leads to increase immature neutrophils in the meninges. Statistical significance, p>0.05; Student’s t-test.

## Discussion

The results in this report clearly demonstrate that the meninges act as a reservoir of neutrophils, and its manipulation can lead to behavior changes. Depletion of neutrophils leads to a deficit in cognitive flexibility, lower interactions with inanimate objects and higher interactions with unfamiliar mice. This depletion paradigm significantly decreases the neutrophil population within the meninges. Within the meninges, neutrophils are not restricted to a specific meningeal layer. The meningeal neutrophil population is heterogenous and populated partially by neutrophils from the circulation and progenitor cells found within the meninges. Upon depletion, the neutrophil progenitor population reenters the cell cycle to generate more neutrophils. We further show evidence that suggests neutrophils in the meninges migrate to adjacent skull bone marrow which is consistent with neutrophils in other spaces that migrate for degradation. Finally, we show that enriched environment and injury have no significant effect on the number of immature neutrophils within the meninges. However, mild stress does lead to a significant increase in the proliferation of these cells.

Recent reports have demonstrated that neutrophils play a diverse role in tissue homeostasis. In liver injury, neutrophil infiltration is critical for the initiation of tissue repair mechanisms^32,33^. In the spleen, neutrophils activate macrophages in the presence of a Candida *albicans* infection ^30,34^. They play the role of antigen presentation to induce B cell maturation in the follicles of the spleen^35^. They also induce the maturation and activation of T cells by becoming antigen presenting cells in response to interferon gamma and granulocyte-macrophage colony stimulating factor ^36^. In an optic nerve crush model, the presence of immature neutrophils is necessary for the induction of nerve regeneration ^37^. Finally, neutrophils release of IL-17 is critical to the activation and function of Th17 cells ^38,39^. In light of these roles, it is important to understand the role of neutrophil in the CNS and meninges. Questions remain as to whether immature neutrophils and neutrophil proliferation in stress plays a specific role in brain function and behavior.

Our understanding of the role of neutrophils in the meninges is still limited. We now know that immune cells play an important immunological role in the CNS. Indeed, T cell subtypes^17^, B cells^40^, and mast cells^18^ have been linked to regulation of cognitive functions. In the periphery, neutrophils play an immunomodulatory role of other immune cells, as such it is important to determine whether neutrophils have the capacity for a neuromodulatory role of cerebral neurons from the meninges.

Finally, a regenerative role has been observed in the immature neutrophil populations in other parts of the body. The presence of this population within the meninges may present a unique target for pharmaceutical intervention to influence CNS physiology in pathological conditions that are known to have inflammatory underpinnings such as trauma, hemorrhage, encephalitis and multiple sclerosis. If this population can be manipulated to release growth factors ^37^ within the meninges, it could be a therapeutic target to mitigate the effect of injury on the brain parenchyma.

## Conclusions

The results of this report show that a deeper understanding of the innate cell population within the meninges is important. We show that manipulating neutrophil population can cause behavioral changes. Whether these changes are permanent or transient still needs to be determined. We also provide evidence that the meninges contain a source of neutrophils, whether this population plays a direct role in brain homeostasis and meningeal regulation still needs to be determined. As with all studies, there are limitations to the one presented. Two of those limitations are evident in the behavioral and cell migration experiments. The limitation of the behavioral experiment is specific to the depletion paradigm, which was global and not restricted to the meninges. Attempts to deplete neutrophils in the meninges, using intrathecal injections of anti-Ly6G, led to peripheral loss of neutrophils. As for the migration experiment, there are currently no methods to do intravital imaging of the meninges without removing some of the skull. Furthermore, the timing of cell movement, as shown in our experiments makes in vivo imaging not feasible at the moment.

## Acknowledgments

Thanks to Dr. Antoine Louveau for teaching us how to make the bone marrow chimera. Thanks to Dr. Chia-Yi (Alex) Kuan for his input during the interpretation of the data. Mariam Syed for contributions to pilot data.

## Methods and Materials

### Animals

Young (8-12 weeks old) male mice were used in all experiments. All experiments were conducted on the following strains:C57BL/6 wild type mice, LysM-tdTomato (LysM-cre [B6.129P2-Lyz2^tm1(cre)lfo^ X Ai14 [B6; 129S6-Gt(ROSA)26sor^tm14(CAG-tdTomato)Hze^]) mice, and GFP (C57BL/6-Tg (UBC-GFP)30Scha/J) mice. Mice were kept on a 12 hour:12-hour light cycle at room temperature (22-25°C). Food and water were provided *ad libitum*. All experiments were done with the approval of the University of Virginia Animal Care and Use Committee.

### Bone marrow chimera

Bone marrow chimera mice were generated by irradiating the LysM-tdTomato mouse and repopulating the bone marrow with isolated bone marrow from the UBC-GFP mouse. In brief, LysM-tdTomato mice received 2 doses of radiation, first 350 rad and second 950 rad, separated by 48hours. Importantly, the radiation doses were given without head shielding so that skull marrow and meninges are included in the radiation field. Following the second radiation dose, mice received an intravenous injection of 5×10^6^ marrow cells isolated from the GFP-mouse long bones. Mice were then kept for 6 weeks then euthanized and the meninges neutrophil population characterized.

### EDU experiments

To determine division in meningeal neutrophils (and other tissues), mice were injected with the thymidine analog 5-ethynyl-2-deoxyuridine (EdU) at a concentration 0.04 g/kg. Mice received 1 intraperitoneal injection of EdU. Mice were then sacrificed, 1-, 2-, and 3-weeks post injections. The meninges, skull and bone marrow were harvested and analyzed for EdU incorporation in the neutrophil population.

### Anti-Ly6G injections

To induce neutrophil depletion, mice were injected with anti-Ly6G (Clone 1A8; BioXcell, Lebanon, NH; 4mg/kg) or saline intraperitoneally. On the day of harvest, blood and meninges were collected from injected mice and flow cytometry performed was performed (as described below) to determine the composition of neutrophil population within each tissue type.

### Flow cytometry

Flow cytometry was performed on the meninges as previously described ^14^. To summarize, a single cell suspension was generated by digesting dissected meninges using 1 mg/ml DNAse and 1.4 U/ml collagenase in Hanks Balanced Salt Solution with magnesium and calcium. Samples were then incubated in a 37°C water bath, triturated and strained through a 70 μm cell strainer. Samples were then centrifuged and resuspended in DMEM media with 10% BSA and kept on ice until staining. Once single cell suspensions were obtained, samples were blocked with Fc block and incubated with an antibody cocktail. To determine cell viability within each suspension, an aliquot of each sample was incubated with the fixable viability dye (ThermoFisher, Waltham, MA, US; 1:1000) for 30 minutes at 4°C.

For samples of mouse blood, cardiac blood was collected at the time of perfusion. Red blood cells were lysed using the ammonium chloride potassium (ACK) lysis buffer (Fisher Scientific) and centrifuged. The resulting pellet was resuspended in DMEM media with 10% BSA and processed with the same staining cocktail as the meninges.

For samples of mouse bone marrow, bone marrow was flushed from mouse long bone or skull bone using a DMEM media with 10% BSA and an insulin syringe. The solution was then passed through a 70 μm, centrifuged and resuspended. The resulting single cell suspension was then stained with the same antibody cocktail used for the meninges.

#### Image stream

For image stream analysis, cell suspensions were incubated in two antibody cocktails. The extra-cellular antibody cocktail contained Ly6G-PE (BioLegend, San Diego, CA; 1:100), CD11b-efluor 780 (Life Technologies, Carlsbad, CA, US; 1:100), CD45-Pacific blue (Biolegend, San Diego, CA, US; 1:200), and Live/Dead aqua (ThermoFisher Scientific, Waltham, MA; 1:1000) for 30 minutes at 4°C. Cells were then fixed with 0.1% paraformaldehyde solution and permeabilized with 0.5% tween solution in 1x phosphate buffer saline (PBS). Cells were then incubated with intracellular antibody cocktail containing MPO-FITC (BD Biosciences, Franklin Lakes, NJ; 1:100). Following a final rinse, cells were resuspended in a 1xPBS solution containing the DNA dye, DRAQ5 (ThermoFisher Scientific, Waltham, MA; 1:1000). Cell suspensions were then analyzed using an Amnis ImageStream X Mark II cytometer (Cytek Biosciences, Bethesda, MD) and analyzed with the Image Data Exploration and Analysis Software (IDEAS).

#### Spectral flow

For spectral flow analysis, cell suspensions were incubated with the following antibody cocktail: CD45-BV650 (BioLegend, San Diego, CA; 1:200), Ly6G-PECY7 (BioLegend, San Diego, CA; 1:200), CD11B-efluor780 (Invitrogen, Waltham, MA; 1:200), CD133-PE (Invitrogen), CD44-PERCP CY5.5 (Invitrogen, Waltham, MA; 1:200), CD49D-PE dazzle 594 (BioLegend, San Diego, CA; 1:200), CD34-PECY5 (BioLegend, San Diego, CA; 1:200), SCA1-BB515, CXCR4-BV 421 (BD Biosciences, Franklin Lakes, NJ; 1:200), Ckit-APC (BioLegend, San Diego, CA; 1:200), Live/dead aqua (ThermoFisher Scientific, Waltham, MA; 1:1000), NK1.1-BV711 (BD Biosciences, Franklin Lakes, NJ; 1:200), CD3-BV711 (BioLegend, San Diego, CA; 1:200), CD19-BV711 (BD Biosciences, Franklin Lakes, NJ; 1:200), B220-BV 711 (BD Biosciences, Franklin Lakes, NJ; 1:200). Cells were then analyzed with the Aurora Northern Lights Spectral Flow Cytometer and analyzed with the Cytobank software.

#### Cell cycle analysis

Single cell suspensions were incubated with Ly6G-PECy7 (BioLegend, San Diego, CA; 1:200) and fixed with 0.1% paraformaldehyde and resuspended in a 1xPBS solution with DRAQ5 (ThermoFisher Scientific, Waltham, MA; 1:1000). Cells were then analyzed using a Attune NXT (ThermoFisher Scientific, Waltham, MA) flow cytometer and analyzed using Flow Jo software.

#### EDU analysis

Single cell suspensions were first generated using the above-described method. Cells were incubated in a cocktail containing external cell markers, such as CD11B-efluor780 (Invitrogen, Waltham, MA; 1:200), CD45-Pacific blue (Biolegend, San Diego, CA, US; 1:200), Ly6C-PERCP Cy5.5 (ThermoFisher Scientific, Waltham, MA; 1:200), Ly6G-PE (BioLegend, San Diego, CA; 1:200), and Live/Dead aqua (ThermoFisher Scientific, Waltham, MA; 1:1000). Following a rinse, cells were processed using the Click-iT Plus EdU flow cytometry assay kit (ThermoFisher Scientific, Waltham, MA). Data was collected using a Gallios cytometer (Beckman Coulter, Brea, CA, US), and analyzed using FlowJo software (FlowJo, LLC, Ashland, OR, US).

### Behavioral Analysis

#### Barnes maze

To determine whether spatial memory and cognitive flexibility were affected by the loss of neutrophils, mice were run in a Barnes maze. To maintain the neutrophil depletion throughout the experiment, mice were injected every 2 days with anti-Ly6G antibodies intraperitoneally. Barnes maze was based on previous protocol used in our laboratory ^14^. The protocol was expanded to test reversal learning and strength of memory. At the beginning of the task, mice were habituated to the maze for 2 days. Habituation was performed by placing mice in the middle of the maze and leaving them to explore for 3 min. During testing, mice were placed in the middle of the maze and given a total of 150 seconds to find the escape hole. Testing for “normal learning” started the day after habituation ended and continued for 7 consecutive days. At the end of the 7^th^ day, start of the “reversal or flexibility learning”, the goal box was relocated 90 degrees from the original placement. The “flexibility learning” was performed for 5 consecutive days. After which, for the “memory strength”, the goal box was moved to its original location. This phase of the maze lasted for 4 consecutive days. Each trial ended when the mouse entered the escape hole, stayed in the vicinity of the escape hole (defined by a circle of 5 cm diameter around the escape hole) for 5 consecutive seconds, or the 150 second maximum had elapsed. All trials were recorded and analyzed using the tracking software Ethovision 13 (Noldus, Leesburg, VA, US).

#### Puzzle box

Cognitive flexibility was further tested in the neutrophil-depleted mice using a puzzle box. The puzzle box is a relatively new behavioral task used to determine cognition and cognitive flexibility in rodent ^41^. This task requires mice to solve increasingly difficult tasks in order to escape into a dark “safety” box from an open field area. The tasks/obstacles used are an open entry, an entry with an open ramp, an entry with a ramp obstructed by bedding material, and an entry obstructed with a tissue plug. The mice are given 3 minutes (open entry and ramp) or 4 minutes (bedding and tissue plug) to overcome the task to escape.

Before testing started, mice were habituated to the box for two consecutive days. Testing started immediately after the last day of habituation. On the first day of testing, mice were place in the box with the first obstacle (open entry) for one trial, followed by two trials with the second (open ramp) obstacle. On the second day, the first trial was with the second obstacle (open ramp) followed by two trials with the third obstacle (bedding). On the third day of testing, the first trial was with the third obstacle (bedding) followed by two trials with the fourth obstacle (tissue plug). On the fourth and last day of testing, the mice had one trial with the fourth obstacle (tissue plug). All trials were recorded and analyzed using the tracking software Ethovision 13 (Noldus, Leesburg, VA, US).

#### Novel object recognition

To determine whether loss of neutrophils affected recognition memory, a novel object recognition test was performed on each group 24 hours after anti-Ly6G injection. Mice were placed in the open field box with the familiar object and given 30 minutes of exploration time. At the end of the 30 minutes, the novel object was introduced to the environment and mice were given an additional 5 minutes of exploration. Using Ethovision 13 software (Noldus, Leesburg, VA, US), mouse behavior was tracked, and the amount of time spent with each object was recorded.

#### Social interaction

To determine whether social behavior was affected by neutrophil depletion, mice performed a social or direct interaction task 24 hours after anti-Ly6G injection. This social interaction task has been demonstrated to test mice ability to distinguish between familiar and non-familiar mice ^42,43^. Briefly, in this task, mice are placed in a previously explored open field box. A non-familiar (i.e. novel) mouse is then placed in the box. Mice are left to interact for 2 minutes. After which both mice are returned to their home cages. After an intertrial period of 1 hour, the testing mouse is placed back in the same open field box. Followed by the addition of either a new non-familiar mouse or the same mouse used for the first trial (now familiar). These mice are then left to interact for 2 minutes. Using Ethovision 13 software (Noldus, Leesburg, VA, US), mouse behavior was tracked, and the amount of time spent with each mouse in each trial was recorded.

#### Enriched environment

To determine if physiological conditions can alter the proliferation rate of immature neutrophils, mice without neutrophils (24 hours after anti-Ly6G depletion) were placed in an enrichment box (an open field box with toys of various sizes and shapes) and left to explore for 1 hour. At the end of the exploration, mice were euthanized, and their meninges harvested for flow cytometry.

#### Acute stress

To determine the physiological effect of stress on the proliferation of immature neutrophils, mice were place in a restrain apparatus for 1 hour. Followed which the meninges were harvested, and flow cytometry performed.

#### Subarachnoid hemorrhage (SAH)

SAH or sham surgeries were performed as previously reported^14^. Briefly, mice were anesthetized, placed in a prone position, and a 3mm incision made on the back of the neck. A conserved subarachnoid vein was then punctured and allowed to bleed. For the sham experiment, the procedure was the same except the vein was not punctured.

### Statistics

Graphpad Prism 9.3 (Graphpad, La Jolla,CA, US) software was used to analyze all of the data obtained. Student’s t-test or analysis of variance (ANOVA) were used to determine whether differences between treatment groups were statistically significant.

---

**Supplemental figure 1:**
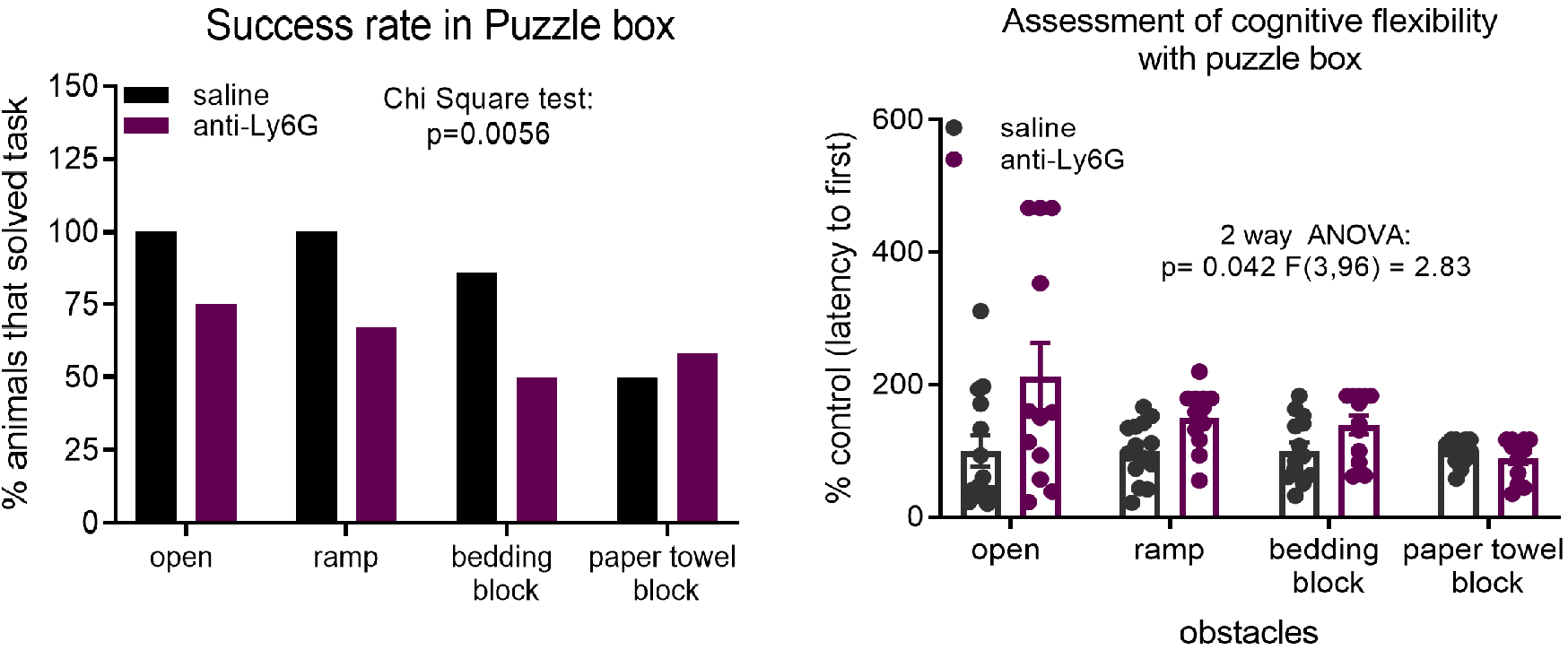
Loss of neutropils alters mouse performance in the puzzle box. Significantly fewer mice, ~25%, did not complete complete the task after the loss of neutrophils. Looking at the latency to escape the box, shows that mice without neutrophils take significantly longer to espace the open field area of the puzzle box.

**Supplemental figure 2:**
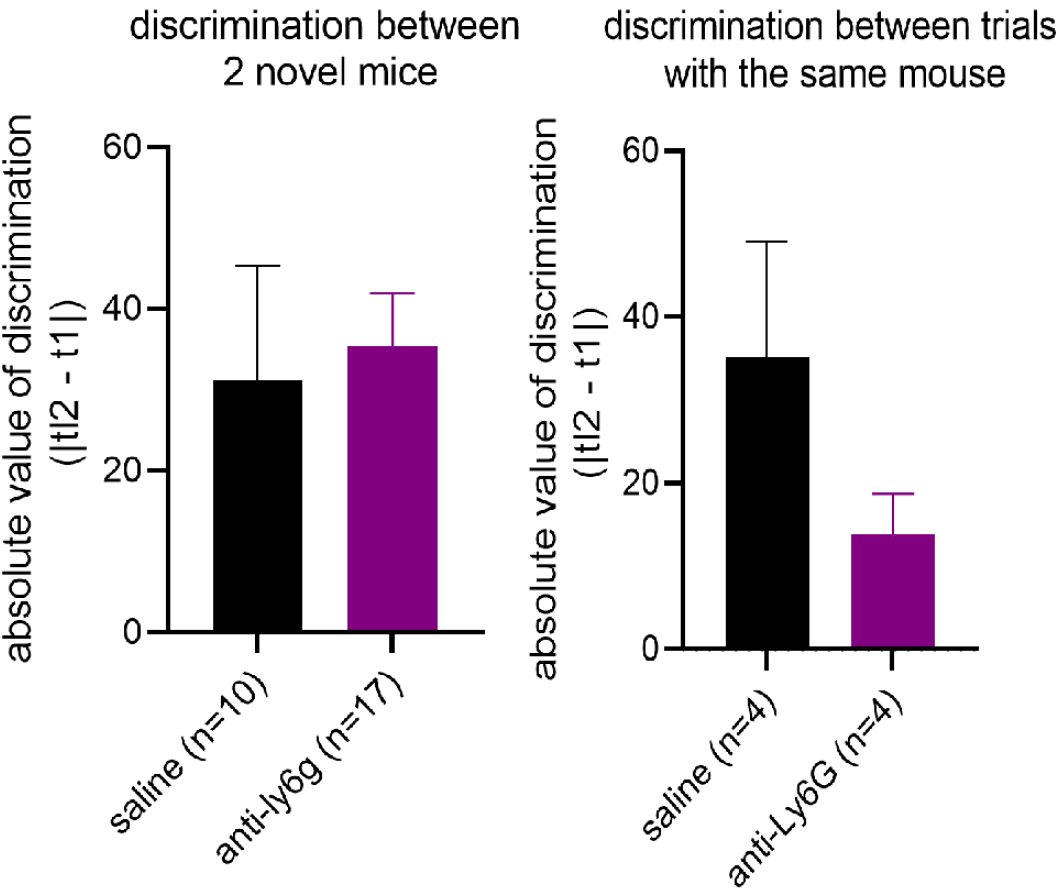
Discrimination index for interaction with either novel mouse once or twice. When introduced to the two different novel mice, the discrimination index is the same between control and neutrophil depleted group. However, if introcuded to the same mouse twice, the saline group dicrimination it enhanced compared to the neutrophil depleted group.

**Supplemental figure 3:**
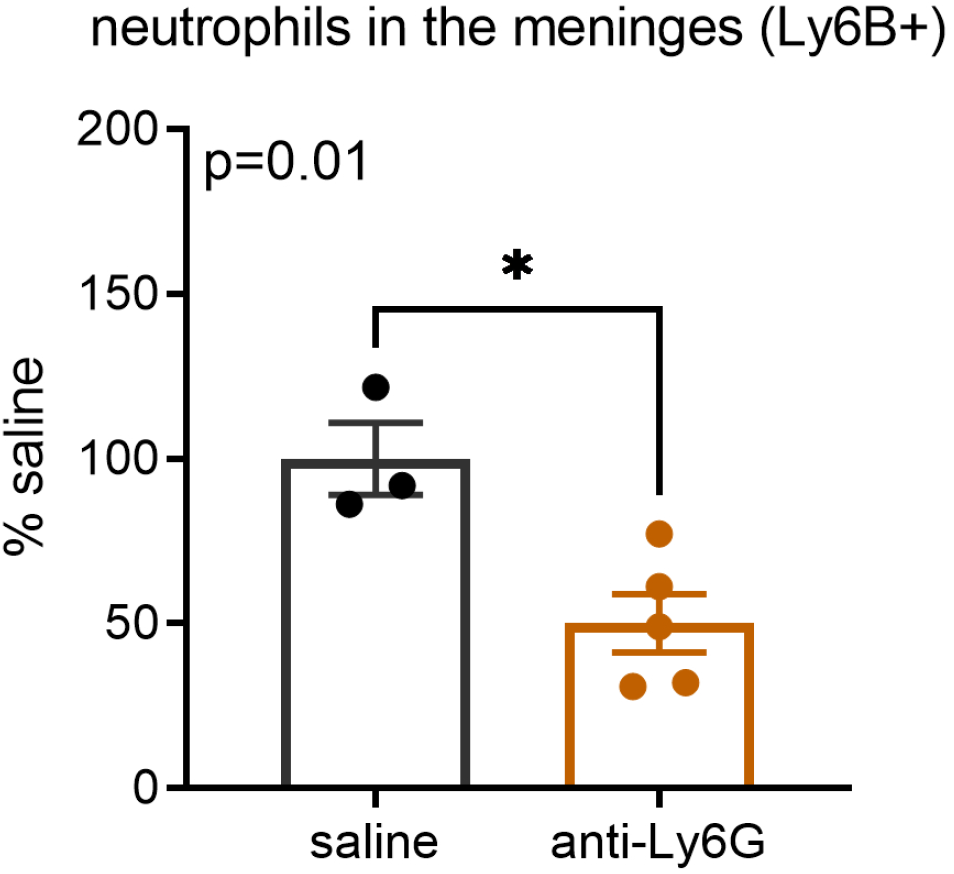
injection of anti-Ly6G antibodies in the peritoneum leads to decrease neutrophils in the meninges.

